# Distinct evolution at TCRα and TCRβ loci in the genus *Mus*

**DOI:** 10.1101/2024.09.05.611428

**Authors:** Moritz Peters, Volker Soltys, Dingwen Su, Yingguang Frank Chan

## Abstract

T cells recognize an immense spectrum of pathogens to initiate immune responses by means of a large repertoire of T cell receptors (TCRs) that arise from somatic rearrangements of *variable*, *diversity* and *joining* gene segments at the TCR loci. These gene segments have emerged from a limited number of ancestral genes through a series of gene duplication events, resulting in a greatly variable number of such genes across different species. Apart from the complete V(D)J gene annotations in the human and mouse reference assemblies, little is known about the structure of TCR loci in other species.

Here, we performed a comprehensive comparison of the TCRα and TCRβ gene segment clusters in mice and three of its closely related sister species. We show that the TCRα *variable* gene cluster is frequently rearranged, leading to deletions and sequence inversions in this region. The resulting complexity of TCR loci severely complicates the assembly of these loci and the annotation of gene segments. By jointly utilizing genomic and transcriptomic data, we show that in *Mus musculus castaneus* the variable gene cluster at the α locus has undergone a recent major locus contraction, leading to the loss of 74 *variable* gene segments. Additionally, we validated the expression of functional variable genes, including atypical ones with inverted orientation relative to other such segments. Disentangling the fine-scale structure of TCR loci in different species can provide valuable insights in the evolution and diversity of TCR repertoires.

## Introduction

T cells are the principal cell type underlying adaptive immunity and perform the remarkable task of distinguishing self from foreign to decide whether or not to initiate an immune response. This pivotal decision depends solely on the recognition of antigens presented by major histocompatibility complexes (MHCs) by the heterodimeric TCR. To cover the enormous space of potential pathogenic antigens, a vast diversity of specific TCR receptors is required. Most T cells express a unique TCR consisting of an α- and a β-chain that arise from somatic rearrangements in a process called V(D)J recombination [1]. Estimates of the diversity generated by this recombination process vary substantially and range from 10^15^ [2] to 10^61^ [3] depending on the mathematical model and the evaluated species. In any case, these theoretical estimates of diversity are several orders of magnitude larger than the observed diversity in any individual (e.g., 2×10^8^ in mice [4] and 1×10^12^ in humans [5]), due to the significantly lower number of total T cells and diversity reduction by selection of specific TCRs during T cell maturation.

The building blocks of TCRs are the variable (V), diversity (D, exclusive to TCRβ) and joining (J) gene segments that are subject to somatic rearrangements by V(D)J recombination. The underlying process requires the precise execution of an ordered series of DNA double-strand breaks that is facilitated by the *Rag1*/*Rag2* recombinase complex [6]. The respective double-strand breaks are repaired by non-homologous end joining (NHEJ) [7] during which random insertions and deletions of nucleotides can occur, which further increases TCR diversity [8]. Recombination signal sequences (RSS) that are interspersed between V(D)J gene segments are targeted by the *Rag1*/*Rag2* complex to initiate recombination. These conserved sequences consist of a heptamer sequence, a spacer sequence with a conserved length of either 12 or 23 base pairs and a conserved nonamer sequence [9]. The 12/23 rule ensures that V(D)J-recombination results in the fusion of a single V to (D) to J segment [10]. The distinctive sequence features of RSS’s are reminiscent of sequences of transposable elements [1]. It is therefore likely that an ancestral version of a TCR gene segment has been invaded by a transposon and subsequently the split gene had to be recombined to encode a functional protein. This hypothesis is supported by the presence of several TCR and BCR related genes in lower chordates that represent potential targets of the initial transposon invasion [11]. V and J gene segment sequences have been categorized into complementarity determining regions (CDR) and framework regions (FR) based on the position of highly conserved amino acid residues in their coding sequence (e.g., cysteine at position 23 and 104 of V genes [12]). The germline-encoded CDR1 and CDR2 regions in the coding sequence of V genes have been shown to modulate TCR-MHC binding affinity [13], while the CDR3 region that comprises the highly diverse junctional region of V(D)J gene segments mainly determines the antigen specificity of the TCR [14, 15].

The distinctive features found in the coding sequence of V(D)J gene segments as well as in their sequence vicinity (e.g., RSS) has allowed their identification from genomic sequences even in the absence of detailed gene annotations [16]. These approaches have revealed that the number of functional V(D)J gene segments varies substantially across taxa and even between closely related species [17–20]. V gene segments are often grouped into families with one to twelve members depending on their sequence similarity of ∼70-100% [21]. TCRβ exclusive usage D gene segments is highly conserved as well as an expansion of the number of J gene segments in the TCRα chain [22]. Similarly, the number of variable gene segments also varies among immunoglobulin heavy chains across different mammals [23]. Successful inference of functional gene segments, however, depends largely on the quality of the respective genome assembly, with complex loci like TCR often representing the worst assembled loci in non-model organisms. This has been emphasized by a recent study that identified V and J genes in the bank vole based on transcriptomic data and identified several additional genes that had not been identified from the genomic sequence [24]. In the same study, most of the identified TCR V and J gene segments were shown to have clear murine orthologs except for three of the identified V genes. In general, significantly less is known about J and D gene segment gene cluster variance, likely because of their relatively short sequence with fewer distinctive features, making it challenging to identify those genes in different genomes. Nonetheless, because both J and D genes contribute mainly to the antigen specificity rather than TCR-MHC binding, the evolution and diversification of their respective loci is particularly interesting in the context of host-pathogen co-evolution.

Locus expansion and contractions of TCR gene segment regions is consistent with the birth-and-death hypothesis of multigene families [25, 26]. This hypothesis is derived from the observation that sequence homogeneity between members of a gene segment cluster within a species is not necessarily higher than to gene segments of a different species [27–29]. It provides the mechanistic basis to explain the evolution of divergent gene segment families, including high frequencies of non-functional and pseudogenized genes following gene duplications and release of functional constraint due to redundancy. While initially evaluated in immunoglobulin and MHC families, subsequent comparative studies of TCR V gene segment families confirmed that sequence identity of homologous families in mice and human exhibit higher similarity than observed between intra-individuals gene families [30]. Later, this view was expanded by showing that divergent Vβ gene segments have been maintained in murine and human genomes for more than 100 million years, strongly indicating that the initial gene duplication events are ancient and predate the split between human and mice [31]. In this context it is important to highlight, that while the diversification of immunoglobulin receptor and TCR loci appears to be driven by similar mechanisms, MHC restriction of TCRs might impose that duplicated gene segments maintain the ability to bind to MHCs to stay functional. In contrast, immunoglobulin gene segments can diversify without such inherent requirements. There now is evidence that four ancestral Vβ and five ancestral Vα gene segments formed the original set of V genes at the root of all mammalian clades. These have since amplified and diverged to different extent in present day mammals [18]. In summary, the birth-and-death hypothesis therefore challenges the classical model of concerted evolution which states that multigene families emerge by inter-locus recombination alongside gene conversion so that all genes within a family cluster evolve in concert and homogenize over time [32].

The murine TCR Vα locus has been subject to one of the most drastic reported locus expansion events in which more than two-thirds of the central locus region has become amplified [21]. Strikingly, this duplication was estimated to have occurred just 4-8 million years ago but has received little attention so far. Today we have access to the high-quality genome assemblies of several common inbred murine strains as well as wild-derived sister species of the most common C57BL/6J laboratory mouse strain [33]. These wild-derived inbred strains share their latest common ancestor about 1-3 million years ago [34, 35] and therefore represent an excellent system to study the evolution of complex traits (reviewed in [36]). Strikingly, the regions with the greatest sequence diversity within the assembled genomes of the various strains relative to the mouse reference genome (GRCm38/mm10) were found to be regions related to immune- or sensory-functions [37]. To date, most comparative studies of adaptive immunity in inbred strains or wild-caught mice are centered around quantifying immune cell populations or measuring differences in immune responses [38, 39]. In contrast, little is known about comparative genomics of TCR gene segment loci despite the fact that those are subject to frequent genomic rearrangements which likely cause significant differences in TCR diversity. Here, we provide a comprehensive comparison of murine TCR loci. By utilizing both, genomic and transcriptomic data, we highlight major rearrangements in the Trav locus of the wild-derived inbred mouse stain CAST/EiJ relative to the mm10 reference and thus emphasize the variability in these loci even in closely related species.

## Results

### The murine TCRα and TCRβ loci in the GRCm38/mm10-based reference

The genomic sequences of the TCR loci have been extensively studied in human and mouse. The gene annotations derived from these studies have been summarized in databases [40] which are now considered to contain all expressed V(D)J gene segments of both species. Here, we now specifically focus on the murine TCR gene segments that are annotated in the IMGT database based on mouse reference genome assembly GRCm38/mm10 (from now on referred to as mm10 assembly). The TCR regions in this database are located in between genes referred to as “locus bornes” (French for milestone) which flank the TCR loci and display an evolutionary conserved gene order across taxa. These can therefore aid the localization of the respective loci. For example, the gene *Dad1*, a 3’ borne. marks the 3’ end of the TCRα loci cluster.

Murine TCR gene segments are found in clusters of varying numbers of gene segments and gene segments within a cluster are further grouped into families bases on their sequence homology and ancestry. The current reference TCRα loci consist of a total of 191 gene segments. These can be further divided into 130 Trav gene segments (including 20 pseudogenes, **Fig. 1A**), 60 Traj gene segments (including 12 pseudogenes, **Fig. 1B**) and a single constant region. All TRCα gene segments are located on chromosome 14 and collectively span about 1.8 Mbp (14C1, 26.94 cM – 27.70 cM). The majority of Trav gene segments consist of two exons with an average span of 556 bp. About two-thirds of the ancestral murine Trav cluster have been triplicated in a recent gene duplication event [21] and all triplicated genes were annotated with a “d” or “n” in their official names to indicate their origin in the ancestral locus configuration. Traj gene segments are all encoded by a single small exon with an average length of 59 bp. The antigen-specificity defining CDR3 region of the TCRα chain consists of the most 3’ bases of a V gene segment and the most 5’ bases of a J gene segment.

**Figure 1:**
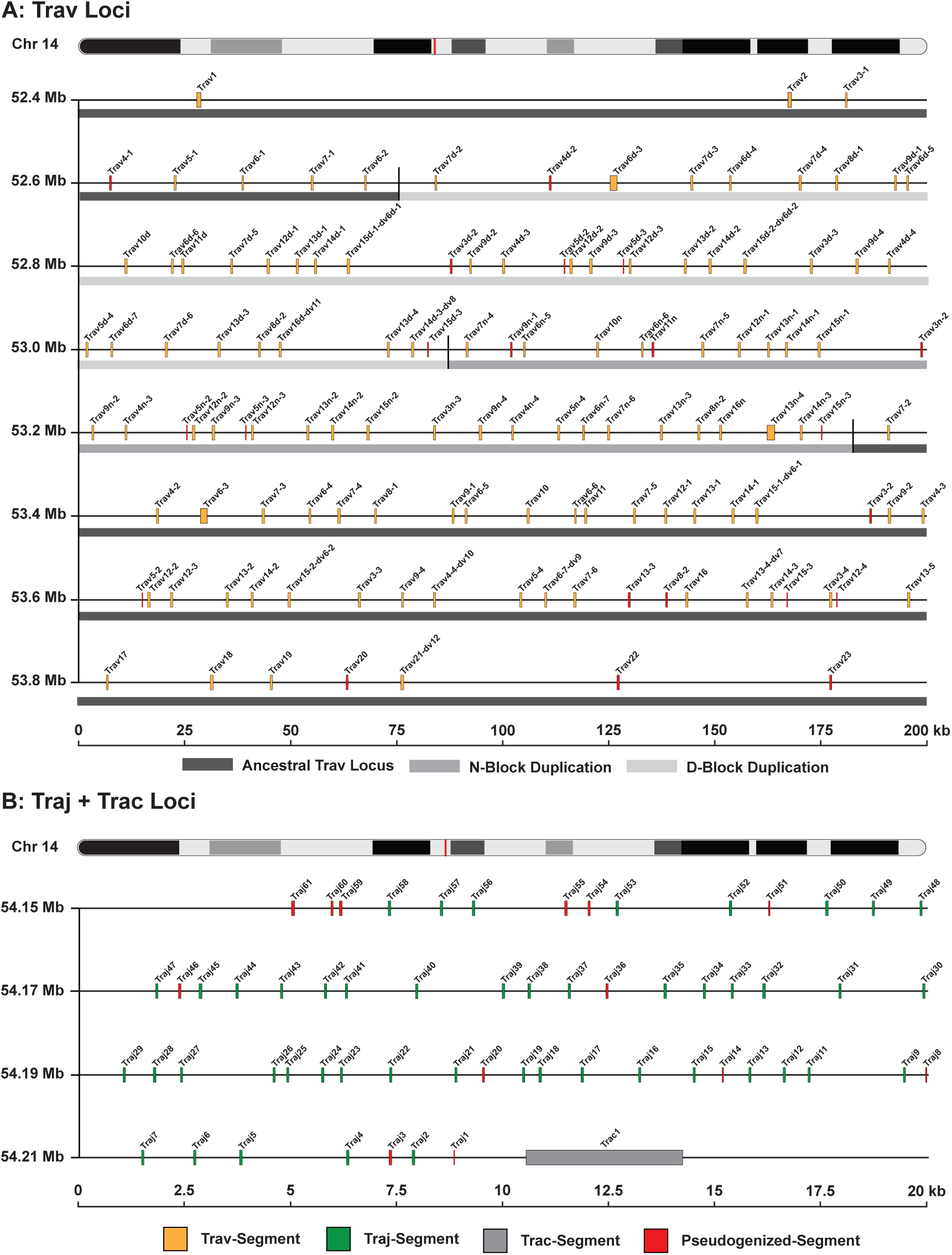
Genomic locations of V/J gene segments in the TCRα loci as annotated in the GRC38/mm10 based IMGT annotation. TCRα *variable* gene segments (130 segments, **A**) are clustered in a 1.6 Mbp genomic region on chromosome 14. A recent gene duplication event led to the triplication of roughly two thirds of the ancestral Trav loci (darkgrey) resulting in the d- (lightgrey) and n-block (middlegrey) Trav segments. Traj and Trac genes (60 Traj and 1 Trac; **B**) are clustered in an 80 kb window upstream of the Trav cluster. Gene segments annotated as pseudogenes are colored in red.

The TCRβ locus spans about 0.8 Mbp and is located on chromosome 6 (6B1, 18.93 cM – 19.71 cM). It consists of 35 Trbv genes (including 13 pseudogenes, **Fig. S1A**), 2 Trbd genes, 2 Trbc genes and 14 Trbj genes (including 2 pseudogenes, **Fig. S1B**). A unique feature of the TCRβ locus is the presence of an inverted V gene segment (*TRBV30* in human and *Trbv31* in mice) at the 3’ end of the locus. Both its position and orientation are conserved in all tetrapods [19]. Across different species the Dβ-Jβ-Cβ clusters are present at varying copy numbers (e.g. 2 in human and mice, 1 in chicken and 3 in swine, [41]). Due to the incorporation of D gene segments, the rearranged TCRβ transcript contains two junctional sites compared to the single junction in the rearranged TCRα transcript.

### The evolution of murine TCR loci

To provide an overview of the TCRα and TCRβ gene segment clusters in other murine species we performed a pairwise sequence alignment of V, D and J gene segments alongside the constant regions in four different inbred mouse lines (129S1/SvlmJ, PWK/PhJ, CAST/EiJ and SPRET/EiJ, from now on referred to as: 129, PWK, CAST and SPRET). For all four species, genome assemblies are made available by the Mouse Genome Project [33]. The dotplot of the local alignments of the Trav cluster confirmed the previously reported locus expansion by triplication of the central region of the cluster in 129, PWK and SPRET (**Fig. 2A**). Strikingly, this local sequence triplication was not observed in the CAST genome assembly and the entire cluster was contracted to a size of about 0.86 Mbp. Apart from this obvious size difference, we also observed local sequence inversions in the Trav cluster which were most frequent in the SPRET assembly relative to the mm10 assembly (**Fig. 2A bottom**). We frequently observed gaps in local assemblies in the genomes of all four mouse strains within the Trav clusters, which in part reflects the complexity of these loci. As a first approach to transfer the reference annotations, we performed a sequence liftover, complemented with a six-frame translation BLAT to identify the chromosomal locations of annotated mm10 V(D)J genes in the assemblies of the four murine inbred lab species (**Table 1**).

**Figure 2:**
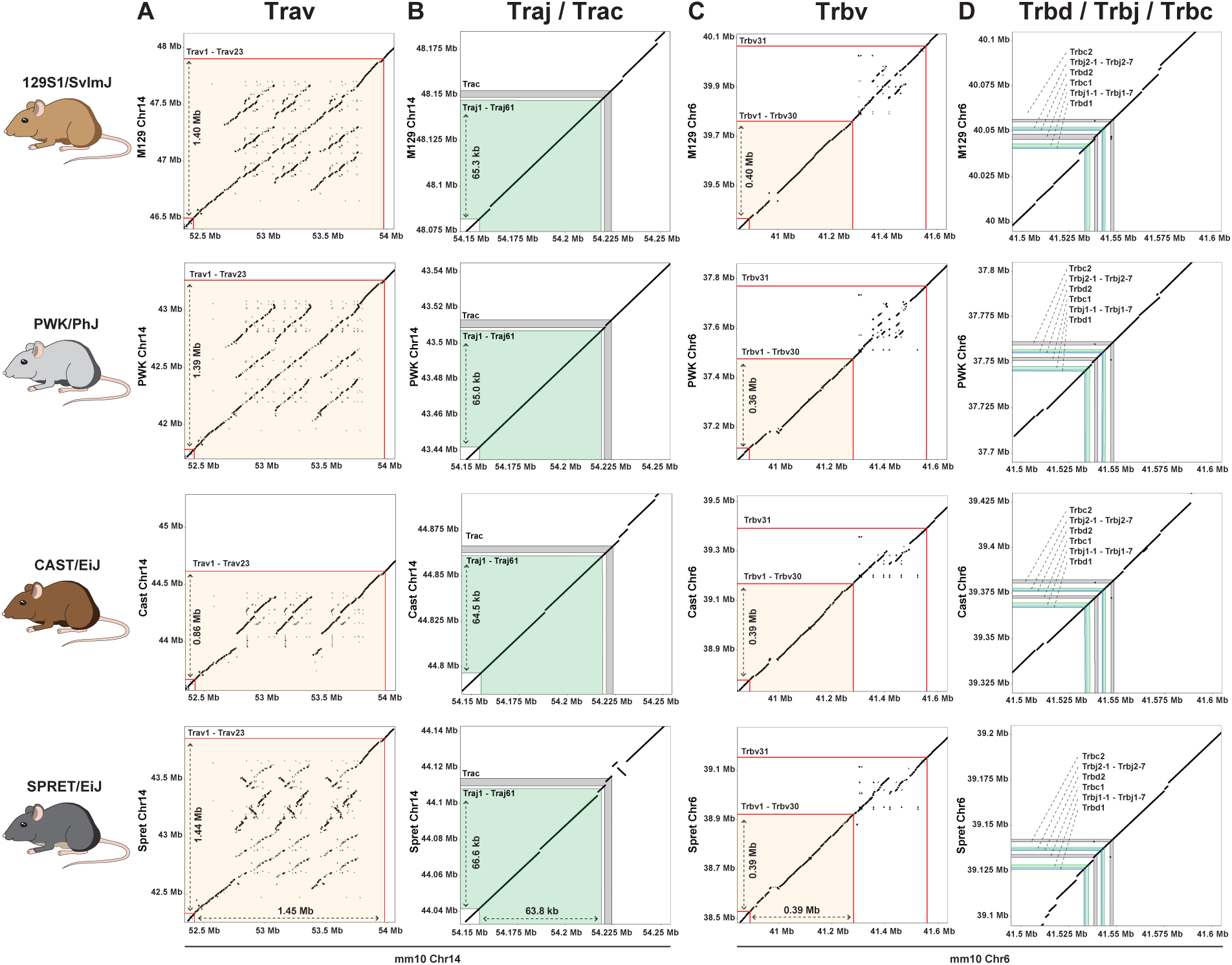
Pairwise alignment dotplots of the TCR genomic regions of four inbred laboratory mouse strains to the respective regions in the murine mm10 reference sequence. Shaded areas illustrate the genomic region ranging from the most 3’ to the most 5’ gene segments.

**Table 1:** Number of V(D)J gene segments (including pseudogenes) identified in the different inbred mouse strains.

| SEGMENT/STRAIN | MM10 | 129 | PWK | CAST | SPRET |
| --- | --- | --- | --- | --- | --- |
| TRAV | 130 | 80 | 97 | 45 (58) | 75 |
| TRAJ | 60 | 59 | 59 | 59 | 59 |
| TRAC | 1 | 1 | 1 | 1 | 1 |
| TRBV | 35 | 35 | 35 | 34 | 35 |
| TRBJ | 14 | 14 | 14 | 14 | 14 |
| TRBD | 2 | 2 | 2 | 2 | 2 |
| TRBC | 2 | 2 | 2 | 2 | 2 |

We did not observe any major locus rearrangement for Jα and Cα (**Fig. 2B**) as well as Vβ,Dβ,Jβ and Cβ clusters and the respective cluster sizes were highly similar among all strains (**Fig. 2C and 2D**). While the *Trbv31* inversion was found in all four strains, we did not observe any further inverted segments. In the following analysis, we now specifically compare TCR loci of CAST to the mm10 reference to highlight the shortcomings of currently available V(D)J gene segment annotations for such complex loci.

### The TCRα and TCRβ locus in *Mus musculus castaneus*

The majority of currently available V gene prediction tools used for the *de novo* annotation of *variable* gene segments in non-model organisms identify candidates by sequence homology to known genes and/or identification of the highly conserved RSS sequences in the vicinity of gene segments [42, 43]. Inherently, these approaches depend on a gapless assembly of the underlying loci, which is often unavailable due to the high complexity V regions.

For CAST we identified a total of 79 full-length *variable* gene segments mapped to the CAST Trav (45) and Trbv (34) region (chromosome 14: 43.65 – 44.65 Mbp, **Fig. 3A** for Trav and chromosome 6: 38.75 – 39.40 Mbp, **Fig 3B** for Trbv). We showed that a large deletion led to the loss of a total of 74 Trav gene segments in CAST relative to the mm10 locus. The deleted Trav segments largely overlap with the triplicated segments in the recent murine Trav triplication event dated back to about 4 - 8 million years ago. Close inspection of Trav sequences alignments against its possible homologs revealed that some of the Trav gene segments in CAST exhibit greater similarity to the corresponding segment in the expanded D-block cluster than to the respective ancestral gene segment. For example, the CAST *Trav6(d)-4* gene showed 100% sequence identity with the mm10 *Trav6d-4* but only 97.5% sequence identity with ancestral *Trav6-4*. Pairwise sequence homology comparison in the remaining Trav gene segments revealed that the deletion junctions are likely located in between the CAST Trav gene segments *Trav7d-4* and *Trav8-1*. We therefore showed, that the “d-“ and “n-blocks” in the Trav locus were present in the ancestors to CAST and thus the present-day CAST Trav locus has undergone a secondary locus contraction, leading to the loss of the majority of the Trav segments in those blocks. All CAST Trav gene segments were subsequently annotated based on the gene segments showing the closest sequence homology in the mm10 reference. Taking into account the latest common ancestor [44] of 129 (*Mus musculus domesticus*), PWK (*Mus musculus musculus*) and CAST (*Mus musculus castaneus*), this contraction has likely happened less than 500.000 years ago. In addition, eleven Trav segments outside the major deletions could not be identified by lifting the genomic coordinates from mm10 to the CAST assembly. For 9 of those Trav segments (*Trav3-1*, *Trav13-1*, *Trav14-1*, *Trav12-2*, *Trav3-3*, *Trav13-3*, *Trav14-3*, *Trav3-4* and *Trav13-5*) we identified corresponding sequence fragments that were terminated by gaps in the CAST genome assembly. Interestingly, the identified Trav3-1 fragment, consisting of just the first exon, showed an inverted sequence orientation (**Fig. 3B**) relative to the homologous sequence in mm10. The two remaining Trav gene segments, *Trav6d-3* and *Trav16*, were not found in the CAST Trav locus because of local sequence deletions (**Fig. 3C**).

**Figure 3:**
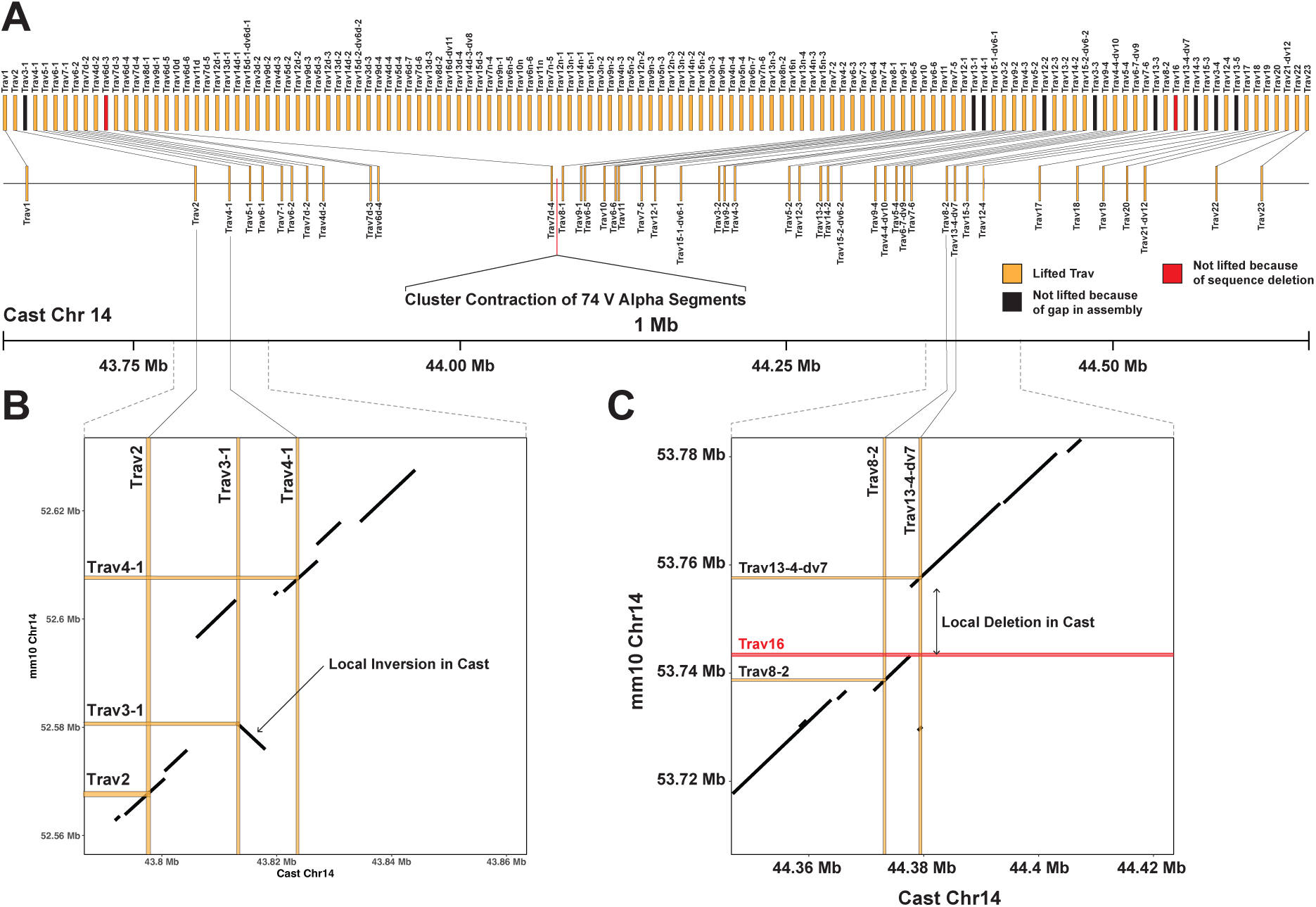
Comparison of the mm10 and CAST Trav gene segment loci. **(A)** Connected segments indicate full-length Trav genes that were lifted to the CAST genome and confirmed in a six-frame translation BLAT (45 in total). Trav gene segments that were not lifted because of gaps in the assembly (black) or are deleted in the CAST genomic sequence (red) lack a connecting line to their mm10 ortholog. **(B)** Zoom-in on the sequence surrounding *Trav3-1* indicates a local sequence inversion in the CAST assembly. **(C)** Zoom-in on the sequence surrounding *Trav16* indicates a local sequence deletion in CAST.

Except for *Trbv9*, an ortholog of all 35 mm10 Trbv segments was successfully identified in the CAST Trbv locus. This was also true for all J gene segments in across both loci (60 Traj and 14 Trbj gene segments; data not shown).

### Gene segment usage validation by gene expression analysis

To validate our gene segment annotation and their usage in the TCR repertoire, we extracted CD8^+^ T cells from the spleen of a 10-week-old male CAST mouse (**see methods**). We then generated TCR repertoire sequencing libraries using the Chromium Next GEM Single Cell 5’ Kit. Critically, this kit utilizes a template-switch based library generation approach, such that it can recover the complete repertoire of expressed TCRs, regardless of the actual recombined 5’ gene segment. Next, we assembled full-length TCR sequences derived from sequencing reads that shared an identical cell barcode and were able to recover a total of 4535 unique Trav and 5389 unique Trbv transcripts (**see methods**). To identify V gene segment alleles, we then collapsed transcripts with identical framework region sequences. To distinguish sequencing and PCR errors from rare alleles, we required each candidate allelic variant to be observed with at least two unique CDR3 sequences. The resulting set of V gene alleles was intersected with the 45 Trav and 34 Trbv sequences generated by direct liftover of mm10 V gene segment coordinates to the CAST genome assembly. For the Trav cluster we identified all 9 full-length coding sequences for the V gene segments that were not identified from the liftover approach, presumably because of assembly gaps in the CAST genome (see previous section). Accordingly, the final set of sequences consisted of 54 Trav gene segments. For all other gene segment loci (Jα, Vβ, Dβ, Jβ) no additional sequences were identified in the transcriptomic data. Next we assembled a full TCRα and TCRβ V(D)J reference library using the *buildLibrary* function of the MiXCR software toolkit [45]. We then used this custom species-specific library to map the sequencing reads of the CAST TCR library, resulting in 86.36% of successfully aligned reads with a V-J spanning clonotype. Next, we analyzed the V(D)J gene segment usage frequencies across all T cells to validate the expression of the identified set of gene segments (**Fig. 4A** and **Fig. 4B**). For Trav genes, we validated the expression of 42 of the 54 V gene segments included in the CAST specific V(D)J reference. The 12 Trav genes that were not expressed consisted of the 9 Trav genes which are annotated as pseudogenes in the mm10-based IMGT reference as well as *Trav13-4-dv7*, *Trav13-5* and *Trav18*, for which we have previously validated the absence of expression in C57BL/6J mice (*Peters et al., 2024, unpublished*). In TCRα chains we observed a prominent pattern of preferential pairing of distal Vα segments with proximal Jα segments and vice versa. This pattern has been described before [46] and provides further evidence for the correct annotation of the underlying gene segments in the TCRα locus. For Traj genes we validated the expression of 44 of the 60 Traj genes included in the respective reference. All of the unexpressed Traj genes are annotated as pseudogenes or ORFs in the mm10-based IMGT reference, for which we have validated the absence of their expression in C57BL/6J (*Peters et al., 2024, unpublished*). In summary, we were able to confirm the expression of 42 Trav genes and 44 Traj genes in CAST mice.

**Figure 4:**
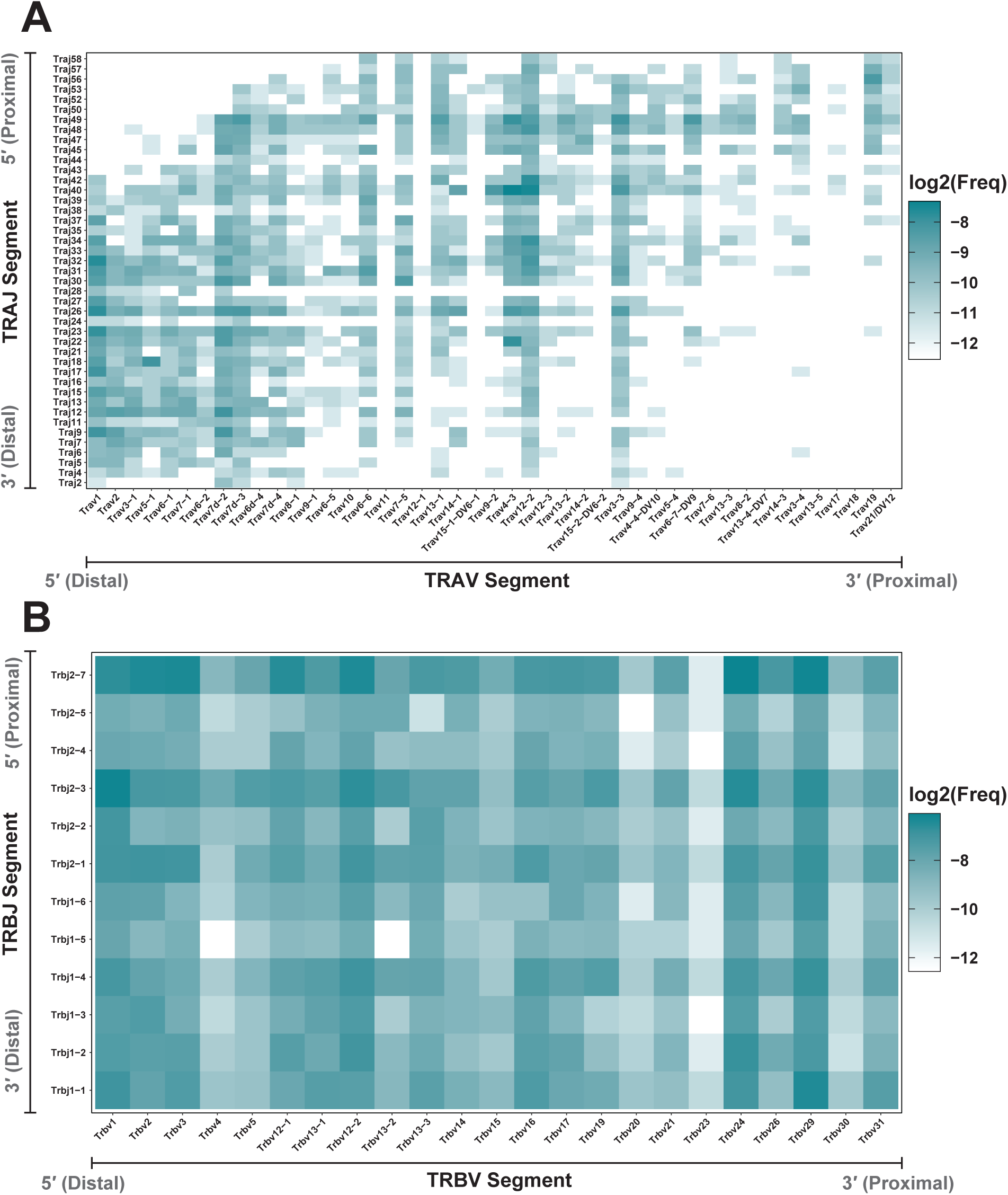
V-J gene segment usage frequency (log2) in the CAST CD8^+^ T cell TCR repertoire. Individual V(D)J gene segments are represented in their chromosomal order for the TCRα (A) and TCRβ (B) chain. In the TCRα chain distal V segments are more frequently paired with proximal J segments and vice versa.

In the TCRβ chains we observed the expression of all 22 Trbv genes that are annotated as functional Trbv genes in the IMGT reference as well as expression of *Trbv21* which is annotated as ORF. We can also confirm the absence of *Trbv24* expression in C57BL/6 (*Peters et al., 2024, unpublished*) and showed that this was caused by a SNP that introduces a premature stop codon (p.Y109X) at the 3’ end of the FR3 region. Critically, in the CAST *Trbv24* sequence this amino acid change was not observed, leading to the frequent utilization of this gene in expressed TCRβ chains. We also observed the expression of all Trbj gene segments that are annotated as functional genes in the IMGT reference as well as expression of *Trbj1-6* which is annotated as ORF. In summary, we were able to confirm the expression of 23 Trbv genes and 12 Trbj genes in CAST mice.

## Discussion

Diversification of immune receptors by somatic recombination is a key feature of the adaptive immune system. The number of available gene segments that are rearranged during V(D)J recombination to generate functional receptors varies significantly in the TCRα and TCRβ chains of different species. These differences are caused by a high frequency of rearrangements in the germline sequence of the underlying gene segment loci, which can lead to heritable locus expansions and contractions. Duplicated gene segments share extensive sequence homology and are therefore grouped into gene segment families. The murine Trav cluster has undergone a recent expansion that resulted in two duplicated blocks (the “d” and “n” blocks) which contain about two-thirds of the ancestral Trav gene segments. In this study we provide evidence for an even more recent rearrangement of the Trav cluster that has led to a major locus contraction in *Mus musculus castaneus* including the loss of 74 Trav gene segments relative to the other sub-species of *Mus musculus* (e.g. *Mus musculus musculus* and *Mus musculus domesticus*). Based on their latest common ancestor, this locus contraction is likely to have occurred less than 500,000 years ago. The frequent sequence duplications leading to highly homologous gene segment family members severely complicates the high-quality assembly of the Trav cluster in reference genomes. At those genomic regions we observed large gaps in the most recent genome assemblies of the four analyzed inbred mouse strains and showed that those gaps often overlap with the predicted location of Trav gene segments. Consequently, V gene segment inference from genomic sequences is prone to yield incomplete gene segment repertoires due to the lack of available sequence information. By utilizing transcriptomic data of CAST TCR receptors, we were able to confirm the expression of 9 Trav gene segments that we were unable to infer from the respective genomic sequences. Critically, we also identified a functional Trav gene segment (*Trav3-1*) with inverted sequence orientation. To the best of our knowledge, functional inverted V gene segments have not been reported for the TCRα chain and have previously only been observed in the form of the highly conserved inverted Trbv gene segment (e.g. murine *Trbv31* and human *TRBV30*) at the 3’ end of the TCRβ locus. Based on pairwise sequence alignments, we showed that large sequence inversions are also present in the Trav clusters of other inbred mouse strains (e.g. SPRET) and therefore likely depict a common feature of rearranged TCR loci that can contain functional gene segments.

In contrast, the remaining gene segment clusters (Jα, Vβ, Dβ, Jβ) showed significantly less major sequence rearrangements across the four different inbred mouse strains. Inference of the CDS of those gene segments from genomic sequences resulted in highly similar, and in most cases identical numbers of predicted functional gene segments across all four mouse species/strains. In line with these predictions, we were able to confirm the expression of all Jα, Vβ, Dβ and Jβ gene segments that are annotated as functional in the mm10-based IMGT V(D)J reference database.

Based on our results, we hypothesized the Trav cluster, relative to other gene segment clusters, is evolutionarily favored to undergo frequent rearrangements, leading to cluster expansions and contractions. An excess of Trav gene segments relative to Trbv gene segments is observed in the majority of mammalian species alongside large numbers of Traj gene segments [18]. The temporally ordered generation of TCRs is initiated by TCRβ rearrangements, a process that is stringently controlled by restricted Rag expression and allelic exclusion. A specific checkpoint, termed the β-checkpoint, ensures that only T cells with a functional TCRβ chain progress to the DP maturation stage. In contrast, at the TCRα locus rearrangement is far less stringent with limited allelic exclusion and prolonged Rag expression leading to continuous rearrangements of TCRα chains over an extended period of time. The ability to “test” different TCRα rearrangements in combination with a pre-defined TCRβ during thymic selection, should evolutionarily favor extended periods of Rag expression and a larger set of Trav and Traj gene segments. Additionally, we can show that thymic selection is more likely to reject particular Trbv compared to Trav gene segments, based on their affinity to MHCs of different MHC-haplotypes (*Peters et al., 2024, unpublished*).

It is therefore likely that purifying selection is less strong for TCRα gene segments, and in fact it may be that expansion of V segments may allow more T cells to survive thymic selection, thus contributing to adaptive immunity. Under such a scenario, even severe rearrangements in the germline configuration of these loci can persist. Following this line of arguments, the initially assembled TCRβ chains could be under selective pressure to maintain a baseline TCR functionality (e.g. by showing appropriate MHC affinity), while the TCRα chains exhibit greater flexibility which can facilitate the rapid adaptation to the exposure of varying pathogens.

In this study, we have highlighted the immense diversity of Trav gene segments that can be observed even in closely related species. We showed that utilizing available genomic sequences of model organisms to predict the sequence of these gene segments often yields incomplete repertoires. This is mainly caused by the dynamic changes in the underlying loci including duplications, contractions and inversions which collectively result in frequent assembly gaps for these regions. Because V(D)J gene segments are the building blocks of functional TCRs, variance in available segments should have significant impact on the TCR diversity of an individual. Unraveling the fine-scale structure of TCR loci is therefore crucial to investigate the evolution and functional specifics of adaptive immune systems.

## Materials & Methods

### Mice

All mice were housed in the animal facility of the Friedrich-Miescher Laboratory. Procedures were approved by the (EB 01/21 M). Mice were originally bought from Charles River Laboratories (Sulzfeld, Germany). Spleens were collected from mice aged 9-11 weeks. The following mouse strains were used in the experiments: C57BL/6J (The Jackson Laboratory, Strain #: 000664), CAST/EiJ (The Jackson Laboratory, Strain #: 000928.

### Isolation of CD8a^+^ T-cells

Spleens of euthanized mice were collected and placed on a 40µm cell-strainer. Spleens were then pressed through the strainer using the backside of a syringe plunger. After thorough rising of the cell-strainer using ice-cold PBS, the flow-through was centrifuged at 400xg 4°C for 10 minutes in a swing-bucket centrifuge. Afterwards, supernatant was carefully discarded and the cell pellet was resuspended in 1ml ice-cold PBS + 2% FBS. Isolation of CD8a^+^ T-cells was then done using the “Dynabeads™ FlowComp™ Mouse CD8 Kit” (Invitrogen, 11462D) according to the manufacturer’s instructions.

### Single-cell TCR sequencing library preparation

After isolation of CAST T cells, TCR sequencing libraries were generated using the 10x Genomics Immune Profiling platform (Chromium Next GEM Single Cell 5’ Kit v2) according to the manufacturer’s instructions. T cells were processed in two separate reactions (two wells of a 10x chip), each with 2.500 input cells. V(D)J sequencing libraries were sequenced at 5.000 reads/cell. Sequencing was done on the Nova-seq 6000 platform by Illumina using S4 2×150bp v1.5 kits with the following sequencing-cycle set-up: R1: 150 cycles, i7 index: 10 cycles, i5 index: 10 cycles, R2: 150 cycles.

### TCR sequencing data processing

Raw fastq-files were processed using the *cellranger vd*j software toolkit provided by 10x Genomics with the built-in mm10 based VDJ-reference (GRCm38-ensemble-7.0.0). In this piepeline fragmented reads are combined into full length contigs based on sequence overlaps in reads and matching cellular barcodes. Importantly, high-quality base call polymorphisms relative to the provided V(D)J reference remain unmodified, so that the generated *filtered_contig.fastq* files contain species-specific allelic variants of these gene segments.

### Species-specific V(D)J reference libraries

The generated *filtered_contig.fastq* files were directly passed to the MiXCR alignment step (“*align*”, --species mmu, --preset generic-amplicon --floating-left-alignment-boundary --floating-right-alignment-boundary C --rna) to generate binary vdjca-files. We then used *mixcr exportAlignments* (--dont-impute-germline-on-export -allNFeatures UTR5Begin FR3End) to extract gene-features so that SNPs in candidate-alleles are not modified to match the provided reference. For each candidate V(D)J-allele we then used the extremely unique combination of associated UMI and CDR3 sequences to distinguish low-frequency alleles from alleles generated by sequencing or PCR errors by requiring each allele to be identified with at least two unique CDR3/UMI combinations. The list of identified V,D and J segment alleles was then used to generate a MiXCR compatible reference libraries for each species using the *buildLibrary* function implemented in MiXCR. Since the underlying RNA-based input libraries are generated using template-switching rather than multiplex-PCR, they allow for the discovery of *de novo* V(D)J-segments since template-switch based cDNA libraries do not require previous knowledge of the entire set of gene-segments for amplification.

### Alignment of sequencing reads using MiXCR

Raw fastq-files containing TCR sequencing reads were integrated into a custom MiXCR pipeline (MiXCR version 4.5.0) using the following steps:

1. *mixcr align*

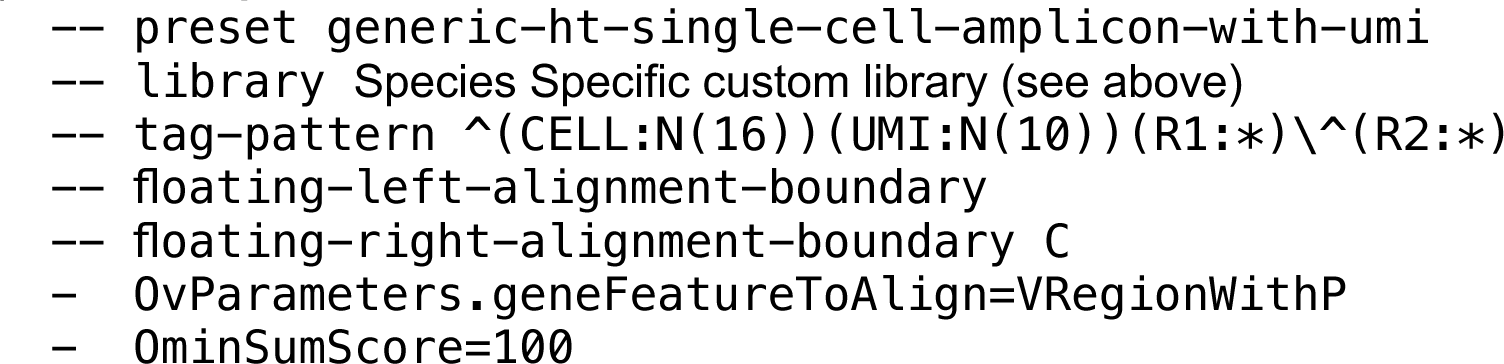
2. *mixcr refineTagsAndSort*
3. *mixcr assemble*

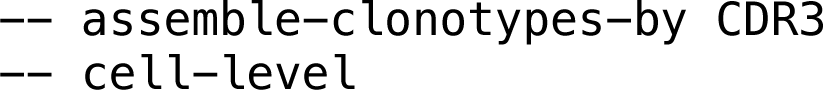

We then used *mixcr exportClones* to extract the required information for all downstream analysis (e.g., cellular barcodes, transcript counts, V(D)J segments, CDR3 amino acid and nucleotide sequence etc.).

### Reference genome assemblies

All assembled murine reference genomes were received from the Ensemble database (release 102). The following reference genomes were used: mus_musculus_129s1svimj, mus_musculus_pwkphj, mus_musculus_casteij, mus_spretus and the standard GRCm38 (mm10) mouse reference genome.

### Pairwise alignment of genomic sequences

We performed a local pairwise alignment of genomic sequences of the TCR loci across all analyzed mice by using minimap2 [47] with the following parameters: “-PD -k19 -w19 -m200 -t48”. The resulting pairwise alignment files (.paf) were then used to plot alignment dotplots using the R package *pafr* [48].

### Liftover of gene coordinates and genome track visualization

Coordinates of the annotated V(D)J gene segments in the GRCm38/mm10 genome were lifted to the genome assembly of the alternative mouse strains using GTF files downloaded from the Ensemble database (e.g. Mus_musculus.GRCm38.102.gtf) and the corresponding “UCSC Chain Files” (e.g. mm10ToGCA_001624445.1.over.chain.gz). The generated GTF files contained the chromosomal locations of the lifted gene segments. These locations were used to generate bed-interval files that were visualized using the Integrative Genomics Viewer [49].

## Acknowledgements

We thank all past and present members of the Chan and Jones laboratory for input into experimental design, helpful discussion and improving the manuscript. We especially thank Felicity Jones for her scientific input throughout the entire study. We thank Sinja Mattes and the remaining team of animal caretakers at the Friedrich-Miescher Laboratory led by Cemal Yilmaz. M.P. and D.S. are supported by an International Max Planck Research School fellowship. Y.F.C is supported by the European Research Council Starting Grant 639096 “HybridMix” and Proof-of-Concept Grant 101069216 “Haplotagging”. The research done in this study is supported by the Max Planck Society.

## Author Contributions

M.P. and Y.F.C. designed the experiments. M.P. performed all experiments with support of V.S.. M.P. and Y.F.C performed the computational analysis. M.P. and Y.F.C wrote the manuscript. V.S., D.S. and Y.F.C. provided support for the experiments and the computational analysis. All authors reviewed the manuscript. Y.F.C. direct the study.

## Declaration of Interest

The authors declare no competing interests.

## Supplement

**Supplementary Figure 1:**
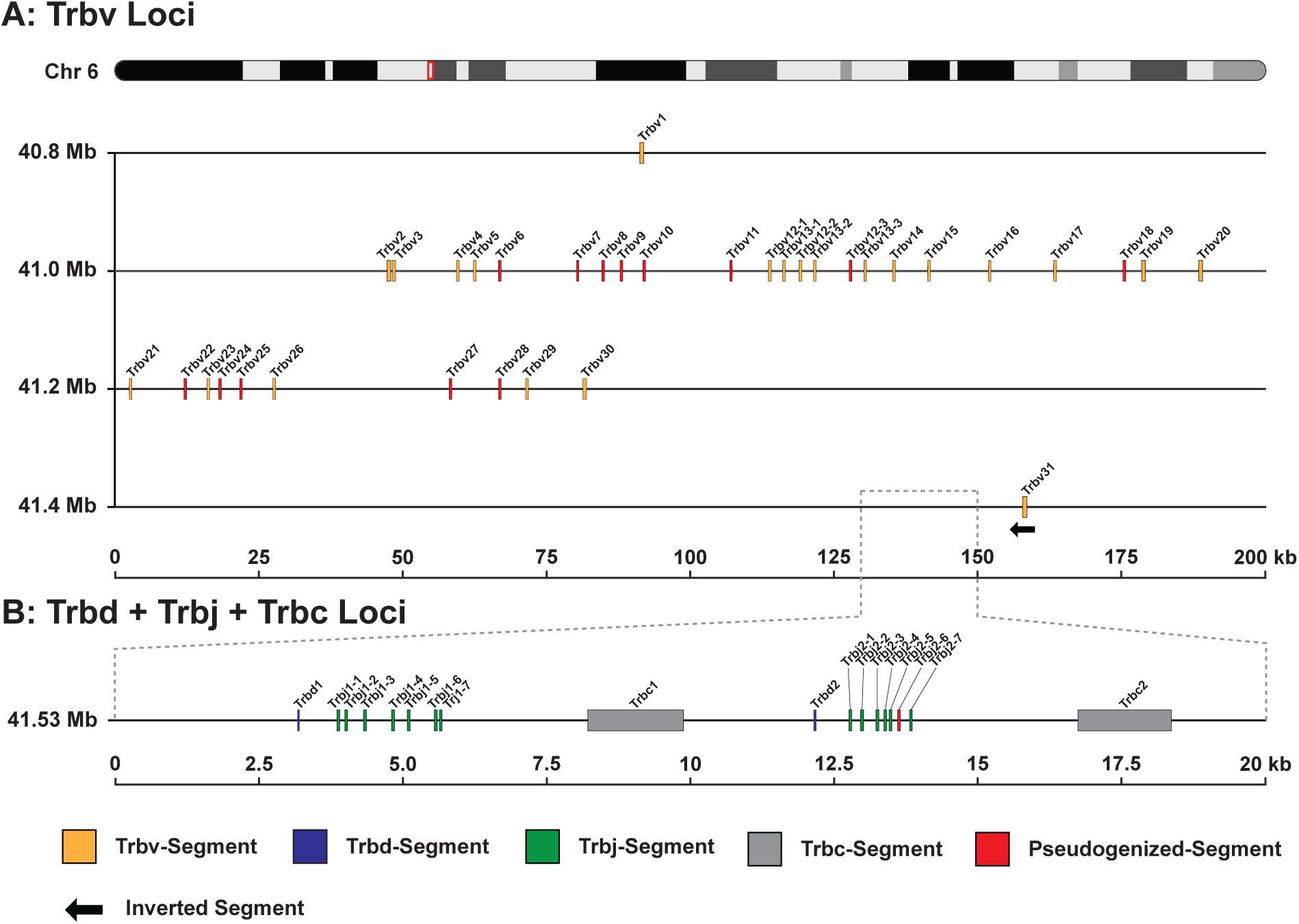
Genomic locations of V/D/J gene segments in the TCRβ loci as annotated in the GRC38/mm10 based IMGT annotation. **(A)** TCRβ variable genes (35 total Trav segments) are located in an 800 kb window on chromosome 6. *Trbv31* is located upstream of the D/J/C clusters in an inverted sequence orientation. **(B)** The D/J/C loci consist of two blocks of a single D gene segment 7 J gene segments and a constant region located upstream of *Trbv1*-*Trbv30* in a 20 kb window.

**Supplementary Figure 2:**
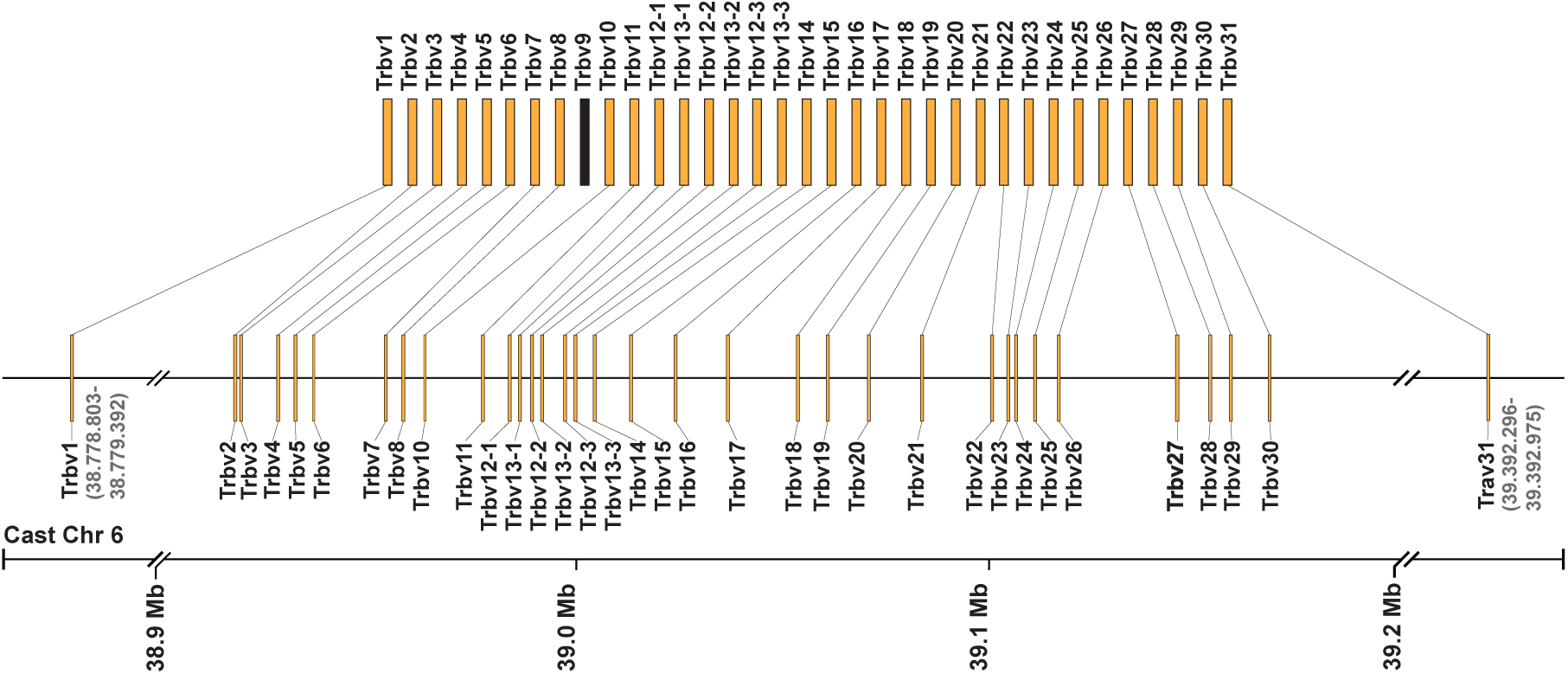
Comparison of the mm10 and CAST Trbv gene segment loci. Trbv gene segments that were lifted to the CAST genome at full-length are connected to their mm10 ortholog. Similar to their location in mm10 *Trbv1* is located downstream and *Trbv31* is located upstream of main Trbv cluster. A homologous sequence of the mm10 *Trbv9* pseudogene could not be identified on chromosome 6 of CAST.

## Notes

### Competing Interest Statement

The authors have declared no competing interest.

### Summary of Updates

The previous submission was found to have rendering errors for in-line figures. This revision fixes the problem. No other change was made.

